# The *Bacillus subtilis yqgC*-*sodA* operon protects magnesium-dependent enzymes by supporting manganese efflux

**DOI:** 10.1101/2024.02.14.580342

**Authors:** Ankita J. Sachla, Vijay Soni, Miguel Piñeros, Yuanchan Luo, Janice J. Im, Kyu Y. Rhee, John D. Helmann

## Abstract

Microbes encounter a myriad of stresses during their life cycle. Dysregulation of metal ion homeostasis is increasingly recognized as a key factor in host-microbe interactions. Bacterial metal ion homeostasis is tightly regulated by dedicated metalloregulators that control uptake, sequestration, trafficking, and efflux. Here, we demonstrate that deletion of the *Bacillus subtilis yqgC-sodA* (YS) complex operon, but not deletion of the individual genes, causes hypersensitivity to manganese (Mn). YqgC is an integral membrane protein of unknown function and SodA is a Mn-dependent superoxide dismutase (MnSOD). The YS strain has reduced expression of two Mn efflux proteins, MneP and MneS, consistent with the observed Mn sensitivity. The YS strain accumulated high levels of Mn, had increased reactive radical species (RRS), and had broad metabolic alterations that can be partially explained by the inhibition of Mg-dependent enzymes. Although the YS operon deletion strain and an efflux-deficient *mneP mneS* double mutant both accumulate Mn and have similar metabolic perturbations they also display phenotypic differences. Several mutations that suppressed Mn intoxication of the *mneP mneS* efflux mutant did not benefit the YS mutant. Further, Mn intoxication in the YS mutant, but not the *mneP mneS* strain, was alleviated by expression of Mg-dependent, chorismate-utilizing enzymes of the <u>m</u>enaquinone, <u>s</u>iderophore, and tryptophan (MST) family. Therefore, despite their phenotypic similarities, the Mn sensitivity in the *mneP mneS* and the *yqgC-sodA* deletion mutants results from distinct enzymatic vulnerabilities.

**Importance:** Bacteria require multiple trace metal ions for survival. Metal homeostasis relies on the tightly regulated expression of metal uptake, storage, and efflux proteins. Metal intoxication occurs when metal homeostasis is perturbed and often results from enzyme mis-metalation. In *Bacillus subtilis*, MnSOD is the most abundant Mn-containing protein and is important for oxidative stress resistance. Here, we report novel roles for MnSOD and a co-regulated membrane protein, YqgC, in Mn homeostasis. Loss of both MnSOD and YqgC (but not the individual mutations) prevents the efficient expression of Mn efflux proteins and leads to a large-scale perturbation of the metabolome due to inhibition of Mg-dependent enzymes, including key chorismate-utilizing MST (menaquinone, siderophore, and tryptophan) family enzymes.

## INTRODUCTION

Metal ion homeostasis relies on a careful balance between metal import, export, and storage mechanisms (1). Bacterial growth may be restricted due to either metal ion limitation or excess, and both mechanisms are important during host-microbe interactions (2, 3). Nutritional immunity refers to the ability of the mammalian host to restrict access to essential nutrient metal ions such as iron (Fe), zinc (Zn), or manganese (Mn) (2). The host may also restrict the growth of intracellular pathogens through metal intoxication (3–5). In response, bacteria induce the expression of metal exporters that contribute to survival in the host (6, 7).

Metal ion efflux systems are regulated in opposition to metal importers. In *Bacillus subtilis,* the MntR metalloregulatory protein binds to Mn to repress the expression of the MntH and MntABCD importers (8). As Mn levels rise further, MntR activates the transcription of the MneP and MneS cation diffusion facilitator (CDF) efflux proteins (9). Although *B. subtilis* normally tolerates up to 1 mM Mn ion, mutants deficient in Mn homeostasis are much more sensitive (9, 10).

Previous work investigated Mn intoxication using an *mneP mneS* (PS) efflux-deficient strain (11). When grown in a minimal medium (MM), the Mn sensitivity of the PS strain can be suppressed by loss of function mutations in *qoxA* (part of the major aerobic respiratory quinol oxidase, QoxABCD) or *mhqR* (repressor of the methylhydroquinone-induced genes) (11). MhqR-regulated proteins function to reduce or degrade quinones and other reactive species (12, 13). Mn intoxication was thereby associated with production of reactive radical species (RRS) due to a dysfunction of the Qox (cytochrome *aa*_3_) quinol oxidase (11).

For reasons not yet understood, genetic suppression of Mn sensitivity is often dependent on the growth medium. For example, in MM with malate as a carbon source, but not with glucose, the Mn sensitivity of the PS mutant was suppressed by an insertion within the *yqgC-sodA* operon (*iTn-sodA*) (11). Conversely, MM with glucose as a carbon source, but not with malate, the Mn sensitivity of the PS mutant was suppressed by the inactivation of *mpfA* (11), which encodes a Mg efflux pump (14). Inactivation of *mpfA* leads to increased intracellular Mg levels, suggesting that Mn intoxication can result from competition with Mg (14). Although Mg is the most abundant divalent cation in cells, it binds macromolecules less tightly than those in the Irving-William series: [Mn(II)<Fe(II)<Co(II)<Ni(II)<Cu(II)>Zn(II)]. Thus, Mg is susceptible to replacement by metals with higher affinities towards protein ligands (including Fe, Mn, and Zn).

Here, we describe the importance of the *yqgC*-*sodA* complex operon in Mn homeostasis. A strain with a deletion of the *yqgC*-*sodA* operon (YS mutant) was defective in the expression of *mneP* and *mneS* and is as sensitive to Mn as a PS efflux-deficient strain. Both strains accumulated high levels of intracellular Mn, displayed high levels of RRS, and had similar alterations in their metabolic profiles. Despite these similarities, the basis for Mn intoxication appeared to be distinct. Several mutations that suppressed Mn sensitivity of the PS mutant had little benefit for the YS strain. Conversely, the YS strain, but not the PS strain, was rescued by elevated expression of chorismate-utilizing, Mg-dependent enzymes of the menaquinone, siderophore, and tryptophan (MST) family. These findings reveal that Mn excess leads to specific metabolic disruptions, and the consequences of these disruptions can vary between phenotypically similar strains.

## Results

### Mutants deleted for the *yqgC*-*sodA* operon (YS) are hypersensitive to Mn

Manganese-dependent superoxide dismutase (MnSOD) is one of the most abundant proteins in the *Bacillus* proteome (15) and is the major Mn-binding protein in the cell (16). The *sodA* gene is transcribed together with the upstream *yqgC* gene as part of a complex, two-gene operon, *yqgC*-*sodA* (Fig. 1a). YqgC encodes an unknown function (DUF456) membrane protein of 160 amino acids. Transcription initiates both upstream of *yqgC* and at two different promoters within the 178 bp *yqgC-sodA* intergenic region (17), with the upstream promoter associated with the production of a 128 nt long 5’-untranslated region designated ncr2103 (18) or S936 (17) (Fig. 1a).

**Figure 1.**
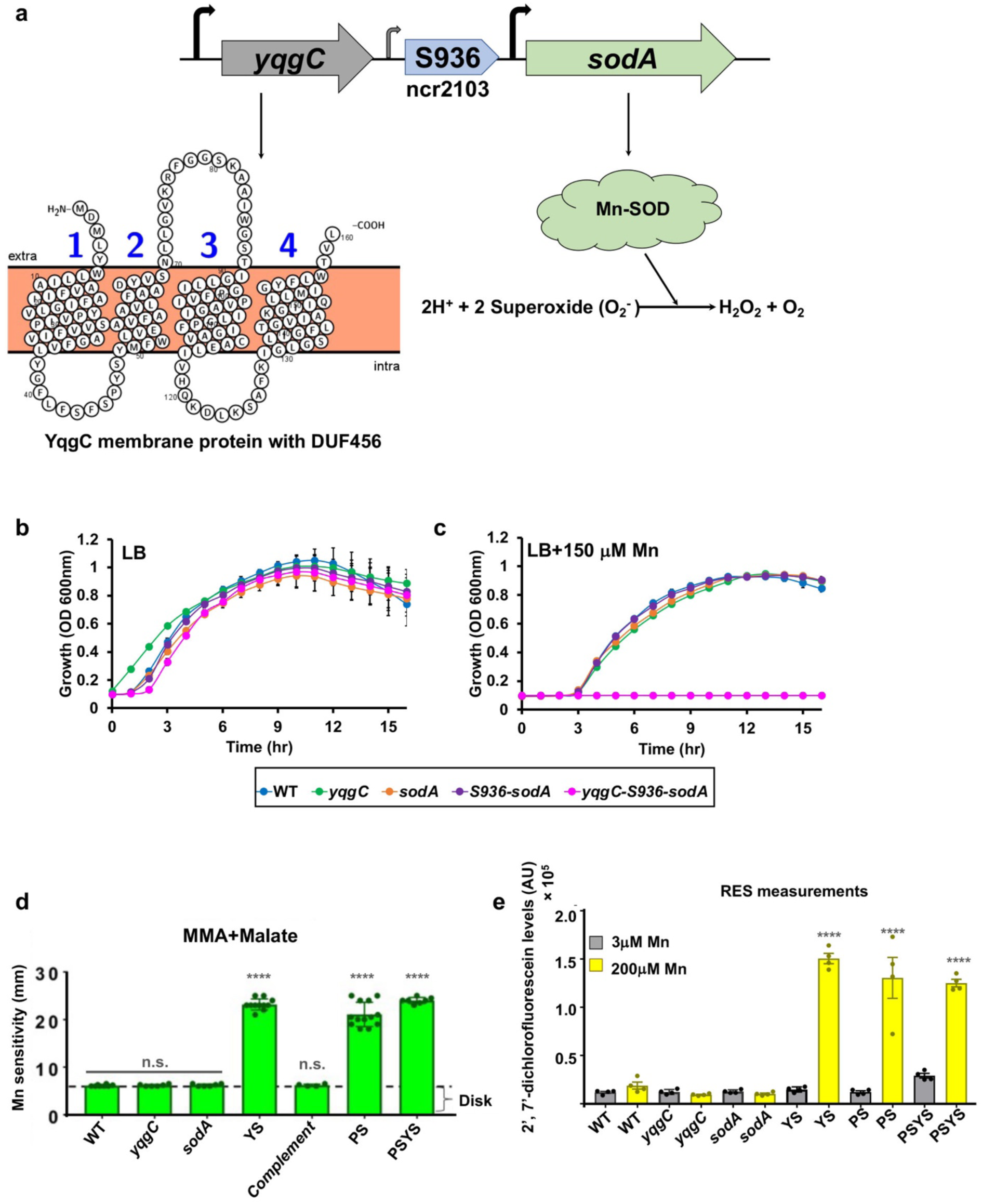
A YS operon deletion strain is hyper-sensitive to Mn. **a)** The *yqgC-S936-sodA* (YS) complex operon encodes (i) YqgC, a multipass transmembrane protein (160 amino acids) with a DUF456 domain (illustrated using Protter (47)), (ii) a 178 bp intergenic region with an upstream σ^B^ promoter (small arrow) that yields a 5’ untranslated region (annotated previously as a 96 nt small RNA (S936) or as non-coding region ncr2103 (18)) and a downstream σ^A^-dependent promoter (larger arrow), and (iii) SodA, a cytosolic MnSOD. Growth characteristics in **b)** LB and **c)** LB supplemented with 150 μM Mn. Time resolved changes in the optical density at 600 nm were monitored periodically for 24 hr under aerobic conditions at 37 °C in a plate reader and were plotted for various strains. **d)** disk diffusion assay was used to assess Mn sensitivity in various strains by monitoring diameter of clearance around a Mn saturated disk (10 μl of 100 mM Mn) on MM-malate agar (MMA) plates supplemented with 0.8 % malate (n=13) **** is adjusted *p* < 0.0001 using a one-way ANOVA test with Tukey’s post hoc analysis; n.s. is the difference of non-significant value. **e)** aerobically grown cells were further exposed to minimal media malate containing either permissive (3 μM) or non-permissible (200 μM) Mn levels and RRS formation was monitored immediately with 2′,7′-dichlorodihydrofluorescein diacetate (DCFDA) under shaking conditions in a plate reader. Plot of fluorescence emission normalized to culture OD is tabulated. RRS production in the presence of 3 or 200 μM Mn were compared using a one-way ANOVA test with Tukey’s post hoc analysis (*n* = 4: \*\*\*\**p* <0.0001).

Previously, we noted that Mn intoxication can lead to an accumulation of reactive radical species (RRS) (11), suggesting that MnSOD may be a protective protein under conditions of Mn intoxication. To explore the role of the *yqgC*-*sodA* operon in Mn homeostasis, we generated in-frame deletions of the *yqgC* and *sodA* coding regions using the BKE collection of mutant strains (19). In these strains, the coding region is replaced by an erythromycin cassette that can be removed using plasmid pDR244 expressing the Cre-Lox recombinase (19). In addition, we used a CRISPR-based mutagenesis approach (20) to delete just the S936 element, the S936-*sodA* region, or the *yqgC-sodA* operon in its entirety (τι*yqgC*-S936-*sodA;* YS). The *yqgC*, *sodA*, and S936-*sodA* deletion strains all grew well in LB medium both with and without amendment with 150 μM Mn (Fig. 1b, 1c). However, the YS operon deletion was unable to grow in LB medium amended with 150 μM Mn (Fig. 1c).

We next quantified Mn sensitivity on MM-malate agar plates (Fig. 1d). The YS strain exhibited a clearance zone equivalent to the Mn-sensitive *mneP mneS* (PS) efflux knockout strain (Fig. 1d). This zone of clearance was not observed in the *yqgC* and *sodA* single mutants. Although both the YS and PS mutants are Mn sensitive, a quadruple mutant (PSYS) was no more sensitive than the YS or PS double mutants (Fig. 1d). Thus, cells lacking the entire YS operon (*yqgC*-*sodA*) are highly Mn sensitive. This high sensitivity was also seen in cells grown in liquid cultures: the MIC for Mn in MM-malate revealed a 400-fold increase in sensitivity in the YS strain (MIC of 5 μM) compared with the wild-type (WT) or single deletions strains (MIC of 2 mM) (Fig. S1). Thus, the presence of either *yqgC* or *sodA* is sufficient to confer resistance to elevated Mn.

### Mn intoxication has similar consequences in YS and PS

Since the YS mutant was as Mn sensitive as the efflux-deficient PS mutant (Fig. 1d), we compared these two strains by monitoring Mn accumulation, production of RRS, and metabolome profiles. When grown in LB medium and challenged with 150 μM Mn, the intracellular level of Mn increased in all strains, but most dramatically in the YS and PS mutant strains (Table 1). After Mn challenge, the YS strain accumulated ∼15x the level of Mn seen in the WT strain, and the YSPS strain accumulated ∼7-fold more (Table 1). In contrast, Mn accumulation was unaffected by the deletion of either *yqgC* or *sodA* (Table 1). In general, elevated levels of Mn were correlated with a decrease in intracellular Mg, but there was little change in Zn levels (Table 2).

**Table 1.**
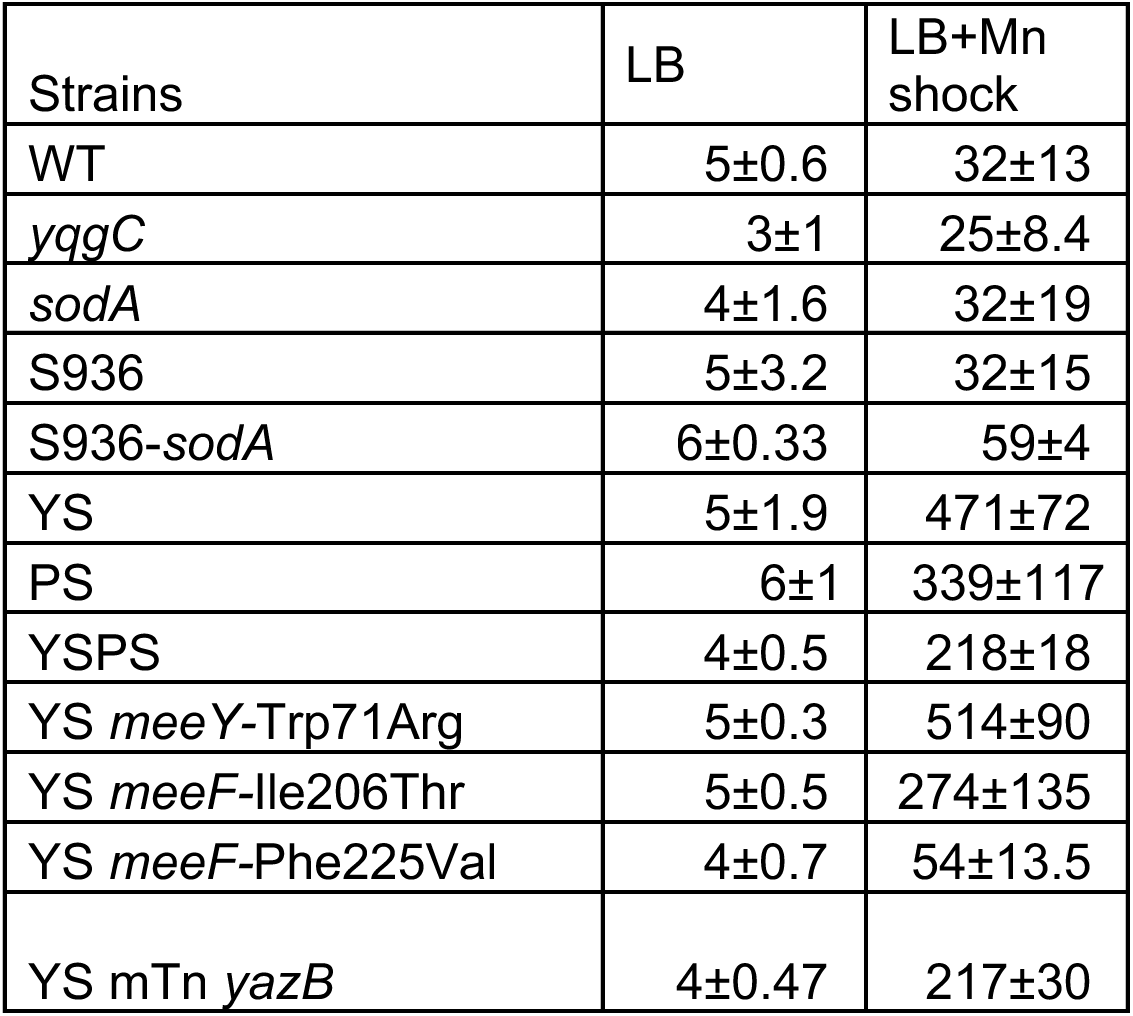
Intracellular Mn levels (μg g^-1^ cells dry weight).

**Table 2:**
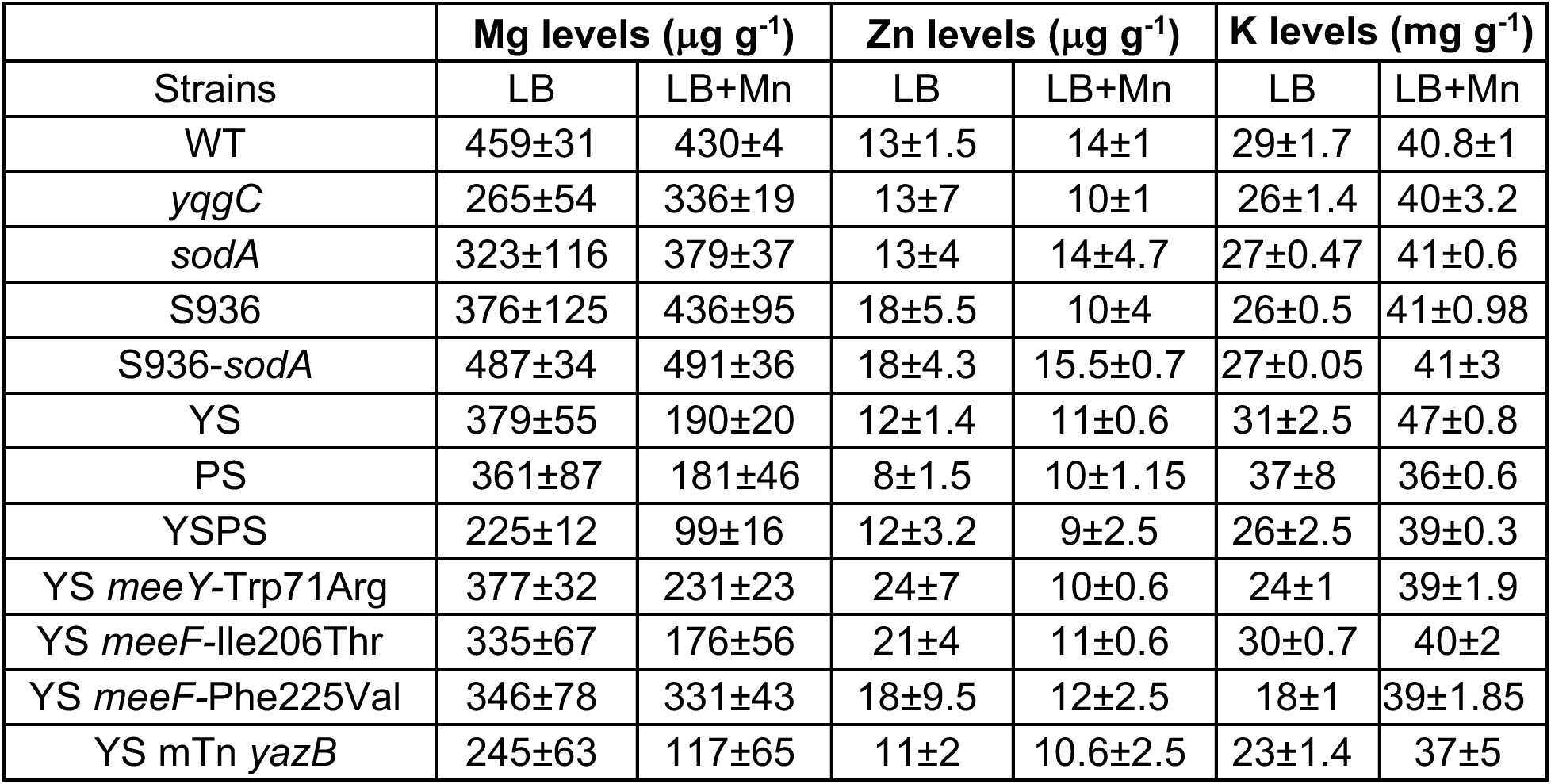
Intracellular Mg, Zn, and K levels.

Previously, the PS efflux deficient strain was found to accumulate RRS when challenged with Mn (11). These studies monitored RRS using the fluorogenic reporter 2′,7′-dichlorodihydrofluorescein diacetate (DCFDA). Intracellular DCFDA is de-acetylated to generate DCF, which is readily oxidized by one-electron oxidizing species, such as hydroxyl radical and other reactive radicals generated by the Fenton reaction (21, 22). Like the PS strain, the YS mutant and the PSYS mutant also accumulated high levels of RRS after Mn challenge (Fig. 1e). We conclude that the YS strain, like the PS strain, accumulates Mn intracellularly and experiences increased stress from RRS.

### Mn sensitivity in the YS strain is correlated with defects in Mn efflux

We hypothesized that the Mn sensitivity of the YS strain may result from a defect in the expression or function of the MneP and MneS efflux pumps. We therefore monitored the levels of *mneP* and *mneS* mRNA using real-time quantitative reverse transcription PCR (qRT-PCR) (Fig. 2a). In the WT background, the three single mutants (*yqgC,* S936, and *sodA*), and the S936-*sodA* mutant, both transcripts were detected and were, as expected, induced by exposure of cells to 150 μM Mn for 30 min. In contrast, in the YS deletion strain the *mneP* and *mneS* transcripts were greatly reduced in abundance and were no longer inducible by Mn (Fig. 2a). These observations suggest that the YS operon deletion precludes efficient expression of *mneP* and *mneS* under these conditions.

**Figure 2.**
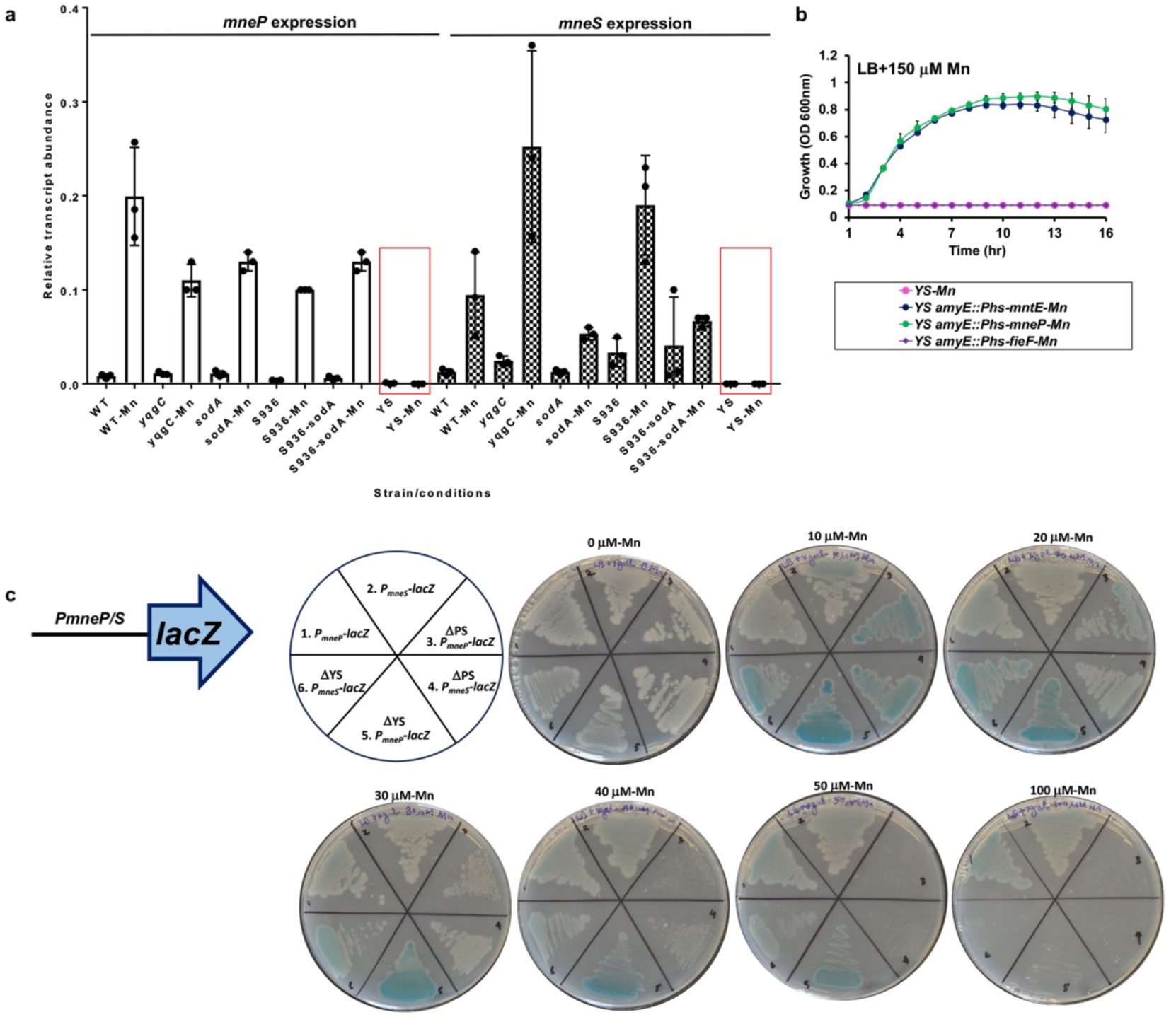
*yqgC*-S936-*sodA* element is needed for *mneP*/*S* expression. **a)** Transcript levels of genes encoding Mn efflux [*mneP* (bars with no fill) *and mneS* (bars with pattern fill)] were monitored in different strains (WT, Δ*yqgC*, Δ*sodA*, Δ*S936*, Δ*S936-sodA,* and YS). Cells were grown in LB till OD_600nm_ = 0.4 and then cells with and without a 150 μM Mn shock for 30 min were harvested and RNA was purified. The expression levels were normalized to the *gyrA* transcript. Values shown are mean ± SD from 3 independent experiments. Red boxes are shown to emphasize the loss of *mneP* and *mneS* transcripts in YS with Mn. **b)** growth characteristics in LB supplemented with 150 μM Mn. Time resolved changes in the optical density at 600 nm were monitored periodically for 24 hr(s) under aerobic conditions at 37 °C. When used P*_hs_-mneP, P_hs_-mntE, and P_hs_-fieF* culture was induced with 0.2 mM of IPTG. **c)** MntR and Mn regulated P*_mneP_* (odd numbers in a pie slice) and P*_mneS_* (even numbers in a pie slice) promoters were fused to β-galactosidase (*lacZ*) gene and were transformed into WT, YS, and PS for comparison of promoter activation. LB agar supplemented with 100 μg/ml of x-gal substrate containing varying levels of Mn were prepared on which mid-log bacterial cultures were streaked for overnight incubation (sectors: 1,2=WT; 3,4=PS and 5,6=YS). The development of blue color is directly proportional to β-galactosidase enzyme activity, which in turn is an indicator for the Mn-based activation of P*_mneP_* and P*_mneS_* by MntR metalloregulator. Note: PS and YS are hypersensitive to Mn and at higher levels show plating defects.

Transcriptional induction of *mneP* and *mneS* by excess Mn is dependent on the MntR transcription factor (9). We hypothesized that the lack of mRNA accumulation for the *mneP* and *mneS* genes might result from a failure of MntR to activate transcription. We therefore tested whether the Mn sensitivity of the YS mutant could be rescued by overexpression of MneP from an IPTG-inducible promoter. Indeed, induction of MneP restored the ability of YS to grow in LB medium amended with 150 μM Mn (Fig. 2b). Similarly, expression of MntE, a *Staphylococcus aureus* CDF (23) that is a homolog of *B. subtilis* MneS (41% identity) and MneP (26% identity), also restored Mn resistance (Fig. 2b). In contrast, expression of the *Escherichia coli* FieF CDF protein (22% identity to MneP, 21% identity to MneS), implicated in Zn, Fe, and Mn efflux (24, 25), did not restore Mn resistance to the *B. subtilis* YS mutant (Fig. 2b). Thus, the Mn sensitivity in the YS mutant strain is due to a failure to properly express Mn efflux pumps (MneP and MneS), rather than an inability of efflux pumps to function properly.

To further test the hypothesis that MntR activation of *mneP* and *mneS* transcription is deficient in the YS strain we monitored the ability of P*_mneP_-lacZ* and P*_mneS_-lacZ* promoter-*lacZ* fusions to be induced by Mn. Surprisingly, amendment with 10 to 20 μM Mn efficiently induced these promoters in the YS strain (Fig. 2c). Expression was similar to that seen in PS strains (Fig. 2c), indicating that activation by MntR is unimpaired in the YS strain. Similarly, the ability of MntR to function as a Mn-dependent repressor, as monitored using a P*_mntH_*-*cat-lacZ* reporter fusion was unimpaired (data not shown). The lack of mRNA accumulation in the YS strain, as seen in LB medium both with and without Mn shock (Fig. 2a), may therefore result from an effect of the YS operon deletion on mRNA synthesis or stability at a step after the MntR-dependent initiation of transcription. The nature of this effect is presently unclear.

### Genetic suppression of Mn sensitivity in the YS mutant

Since the YS operon deletion leads to a reduction in expression of the *mneP* and *mneS* genes, we anticipated that YS and PS mutants might have very similar effects on cell physiology. To test this hypothesis, we asked whether suppressor mutations previously shown to rescue Mn sensitivity of the PS strain (11) also rescue the YS strain (Fig. 3a). Unexpectedly, none of the mutations that improved fitness of the PS mutant strain on MM-glucose (*mhqR, pyk, mpfA, ytfP-opuD*) were able to improve the growth of the YS strain on this medium (Fig. 3b). The mutation that most strongly rescued growth of the PS strain on MM-malate (*qoxA*) had a modest beneficial effect in the YS strain (Fig. 3c). A small beneficial effect was also noted for the *pyk* mutation in YS (Fig. 3c), even though this mutation was not beneficial to the PS strain on this medium (and was selected on MM-glucose (11)). While the basis for these genetic interactions is poorly understood, these differences imply that, despite their phenotypic similarities, the processes that limit growth in the PS and YS strains may be different.

**Figure 3.**
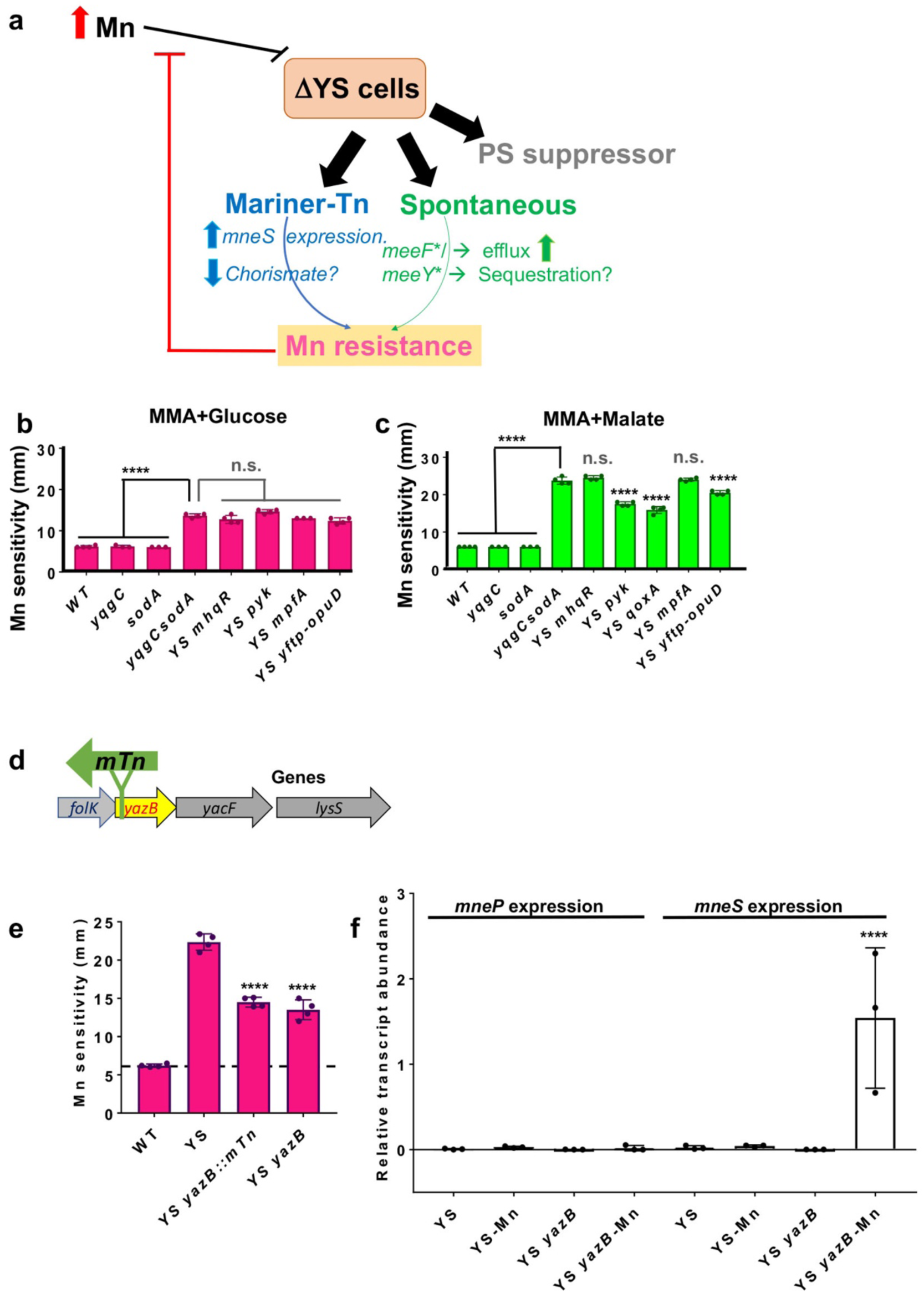
Selection of Mn resistance in YS. **a)** High Mn is toxic to YS cells suggestive of Mn associated sensitivity. To understand the processes failing under such conditions we tested the effect of (i) PS suppressors of Mn intoxication isolated previously (11); Since YS phenocopies PS, with reduced *mneP* and *mneS* gene expression. (ii) newly isolated spontaneous suppressors from the Mn zone of inhibition for YS background, these suppressor mutations were in *yceF* (*meeF)* and *ykoY* (*meeY)* proteins associated with Mn metalation of exoenzymes and (iii) newly isolated *mTn*-generated suppressors in YS on survival of YS under minimal media malate with excess Mn. The PS Mn sensitivity suppressors (*mhqR*, *pyk*, *mpfA*, *qoxA, ytfP-mTn-opuD*) (11) were introduced into YS and Mn sensitivity was determined using a disk diffusion assay at 37 °C. Zone of clearance/sensitivity was measured by enumerating a diameter (mm) after overnight incubation for 18 hr on minimal media agar (MMA) with **b)** glucose or **c)** malate as a sole source of carbon (n=4 and **** is adjusted *p* < 0.0001 using a one-way ANOVA test with Tukey’s post hoc analysis; n.s. is the difference of non-significant value). Note the disk diameter is 6.5 mm. **d)** schematic view of site of mTn insertion in *yazB* is shown for YS Mn resistant mutant leading up to possible upregulation of *mneP/S* genes. **e)** disk diffusion assay was used to assess Mn sensitivity in various strains by monitoring diameter of clearance around Mn saturated disk (10 μl of 100 mM Mn) on MM-malate agar (MMA) plates supplemented with 0.8 % malate (n=4) **** is adjusted *p* < 0.0001 using a one-way ANOVA test with Tukey’s post hoc analysis; n.s. is the difference of non-significant value. **f)** transcript levels of *mneP and mneS* were monitored in YS and YS *yazB* strains. Cells were grown in LB till OD_600nm_ = 0.4 and then cells were incubated with or without 150 μM Mn for 30 min, cells were harvested, and RNA was purified. The expression was normalized to the *gyrA* transcript. Values shown are mean ± SD from 3 independent experiments.

To better understand the effects of Mn on the YS strain we next selected spontaneous mutants that formed colonies in the zone of clearance around Mn disks in zone of inhibition assays. Results from whole genome-resequencing (Supplementary file S1C) revealed that these isolates often carried missense mutations affecting either of two TerC homologs (MeeF, MeeY) implicated in Mn export and exoprotein metalation (26, 27). As expected based on prior work (26), null mutations in *meeF* or *meeY* did not increase Mn resistance in the YS background. Therefore, these missense mutations were recreated by CRISPR-based genome editing to test whether their introduction was sufficient for Mn resistance. In each case, CRISPR-generated strains with only the altered function *meeF*\* or *meeY*\* alleles were Mn resistant. Since TerC homologs are implicated in Mn efflux (26, 27), we used ICP-ES to measure the impact of *meeF\** or *meeY\** alleles on Mn levels. Our ICP-ES results (Table 1) suggest that the *meeF\** allele (Phe225Val) decreased Mn accumulation in the YS cells, likely accounting for the increased Mn resistance. In contrast, *meeY*\* (Trp71Arg) and *meeF\** (Ile206Thr) led to only modest reductions in Mn accumulation (Table 1).

Next, we used mariner transposon (*mTn*) mutagenesis to isolate mutations that could suppress Mn sensitivity of the YS strain (Fig. 3d). We recovered 8 different transposants with insertions in a variety of loci (Supplementary Table S1D), all of which were genetically linked to the observed Mn resistance. We here focused on strain HBYL1110 with an *mTn* insertion after position 87484 in the *Bacillus* reference genome (NC_000964; (28)) within *yazB*. The *yazB* gene is part of a complex operon including genes for folate biosynthesis (*folB*-*folK*-*yazB*-*yacF*-*lysS*) (Fig. 3d). YazB is predicted to be a small (69 amino acid) DNA-binding protein, suggestive of a possible regulatory function. Prior studies of a *yazB* mutant did not reveal changes in the expression of the upstream folate genes (29). We confirmed that an in-frame *yazB* deletion mutant also suppressed Mn sensitivity of the YS strain (Fig 3e).

Since YS strains are defective in expression of *mneP* and *mneS*, we monitored the effect of the *yazB* on these two genes. Remarkably, Mn-dependent induction of *<u>mneS</u>* was restored in the YS *yazB* strain, but expression of *mneP* expression was still non-responsive to Mn stress (Fig 3f). This suggests that the *yazB* suppressor works by restoring the expression of *mneS*, consistent with the conclusion above that YS fails to properly express the MneP or MneS efflux proteins. We attempted to test this hypothesis by constructing a YS *yazB mneS* quadruple mutant, but were repeatedly unable to obtain this strain.

### Mn intoxication elicits similar changes in the metabolome of YS and PS strains

Next, we used metabolomics to monitor the global changes in metabolism upon challenge of WT, *yqgC*, *sodA*, and YS strains grown in LB with and without 150 μM Mn and compared these changes to those seen in the efflux-deficient PS mutant. The *yqgC* and *sodA* single mutants had relatively modest metabolic perturbations after Mn challenge. In contrast, the Mn-sensitive YS and PS strains had similar and wide-ranging changes in cellular metabolism (Fig. S2).

Inspection of the metabolomics results revealed striking effects on chorismate-dependent pathways (Fig. 4a). Specifically, there was an accumulation of chorismate upon Mn-treatment of the *sodA* mutant, as well as the YS and PS strains (Fig 4b). In addition to an elevation of chorismate, the YS and PS strains also displayed a Mn-dependent decrease in downstream metabolites, including the tryptophan precursor anthranilate and the menaquinone precursors 1,4-dihydroxy-2-naphthoic acid (DHNA) and (1R,6R)-6-hydroxy-2-succinyl-cyclohexa-2,4-diene-1-carboxylate (Fig. 4B, S2). These metabolic perturbations led us to hypothesize that Mn might inhibit enzymes of the menaquinone, siderophore, and tryptophan (MST) family (Fig. 4a). *B. subtilis* encodes several MST-family enzymes that are involved in the synthesis of the bacillibactin siderophore (DhbC), folate (PabB), and menaquinone (MenF). MST enzymes bind chorismate, with the carboxylate group coordinated to divalent magnesium (Mg^2+^), and catalyze a variety of addition reactions (with either water or ammonia as nucleophile) that may or may not be coupled to the elimination of pyruvate (30–32). Other Mg-dependent enzymes may also be inhibited under these conditions, including Mg-dependent amidotransferases involved in aromatic amino acid (Trp, Phe, Tyr) biosynthesis. In addition, asparagine levels are greatly reduced in the Mn-challenged YS and PS mutants (Fig. S2), and asparagine synthase is a Mg-dependent amidotransferase (33). Similarly, inhibition of the HisC/AroJ amidotransferase may account for the observed increase in phenylpruvate and decrease in phenylalanine (Fig. S2).

**Figure 4.**
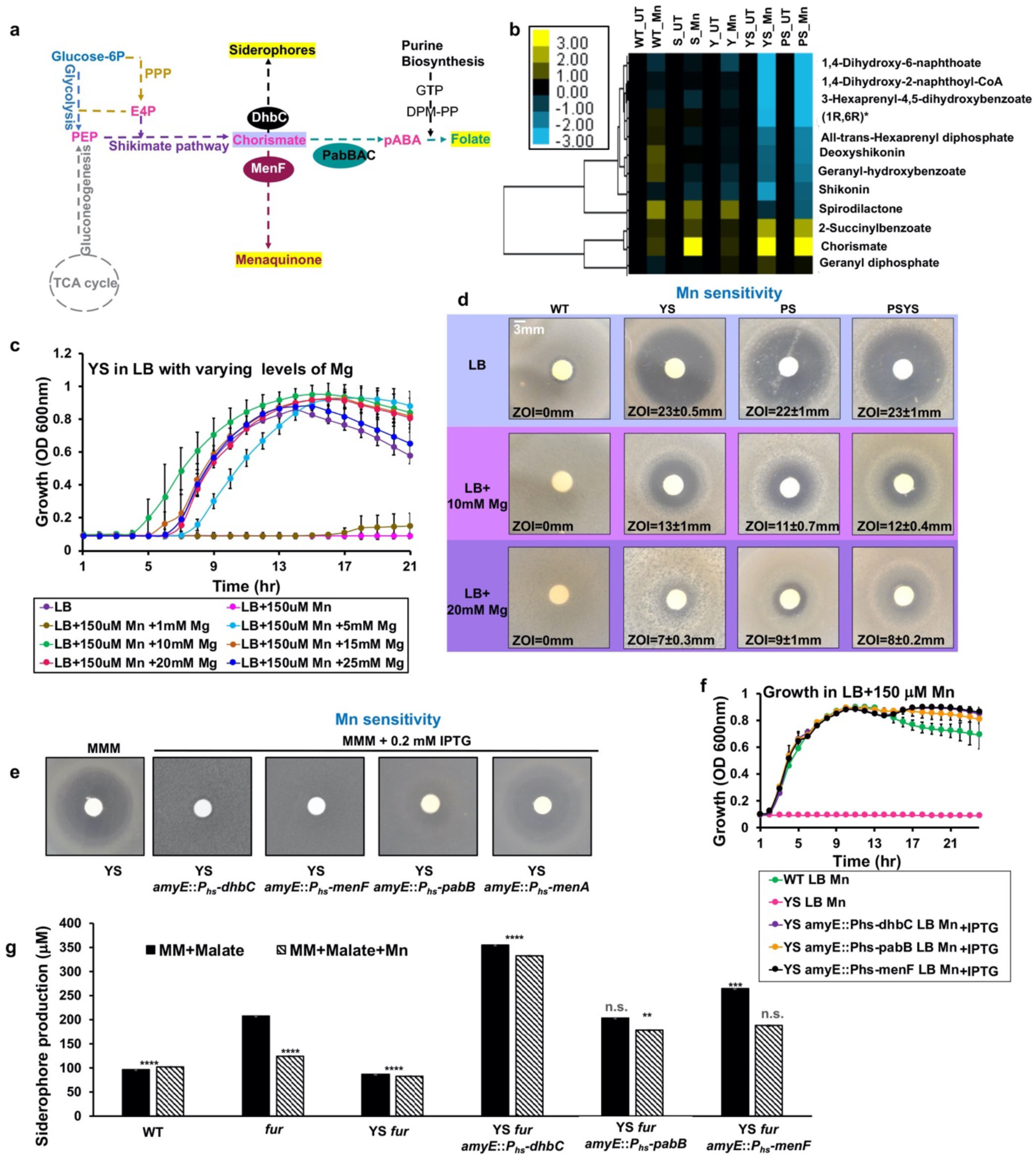
Chorismate-dependent metabolites are affected in the absence of YS. **a)** Schematic representation showing the role of chorismate as a branch-point metabolite. Phosphoenolpyruvate (PEP) is a 3-carbon intermediate generated from primary metabolism (glycolysis and gluconeogenesis shown in blue and grey, respectively) and erythrose-4-phosphate (E4P) is generated by the pentose-phosphate pathway (PPP shown in orange). PEP and E4P are funneled into the shikimate pathway (shown in purple) to generate chorismate. Chorismate serves as a precursor for other metabolites including folate. The *pab* locus encodes Pab proteins (teal circle) that generate the folate intermediate p-amino-benzoic acid (pABA). Other pathways leading to menaquinone, siderophore, and tryptophan are shown. **b)** extracted and measured metabolites from YS and PS strains subjected to Mn using LC-MS/MS are shown. This selective snapshot (see also Fig. S2), from two independent experiments with biological triplicates, highlights the metabolites of the shikimate, folate, and menaquinone pathways. Color bar represents log_2_ fold changes. **c)** growth of the YS strain was tested in LB supplemented with 150 μM Mn or LB supplemented with 150 μM Mn plus varying (1-25 mM) levels of Mg. Time-resolved changes in OD_600nm_ were monitored for 24 hr under aerobic conditions at 37 °C in a plate reader and were plotted for various metal combinations. **d)** Mn sensitivity measured by disk diffusion assay (10 μl of 100 mM Mn) on LB with and without 10 or 20 mM MgCl_2_ supplementation. **e)** Mn sensitivity measured by disk diffusion assay on MM-malate agar (MMA) with and without 0.2 mM IPTG. For enzyme expression (*menF*, *dhbC*, *pabB*, and *menA*), each gene was cloned in pPL82 under P*_hyspank_* promoter (P*_hs_*) and integrated at *amyE* in the YS::*erm* background. Representative images are shown with a consistently magnified view (100X). **f)** growth was tested by inoculating YS and YS overexpressing MST enzymes in LB supplemented with 150 μM Mn, with and without 0.2 mM IPTG. OD_600nm_ was monitored periodically under aerobic conditions at 37 °C in a plate reader. **g)** DHBA-based siderophore yield (absorbance 510 nm/ OD_600nm_) was determined for the indicated strains with and without over-expression of MST enzymes (*menF*, *dhbC*, and *pabB*). Cells were grown overnight in MM-malate with or without 0.2 mM IPTG with 1 μM ferric ammonium citrate, harvested and then resuspended in the same medium with no added iron and growth continued for 5 hrs. Supernatant was used in determination of DHBA levels after colorimetric derivatization with excess FeCl_3_. The level of siderophores/DHBA were extrapolated from DHBA standard curve treated in a similar manner (n=2: the difference in siderophore production among all strains were compared to the *fur* mutant grown in liquid minimal media malate (MMM) without Mn using a one-way ANOVA test with Tukey’s post-hoc analysis where \*\*\*\**p*<0.0001, \*\*\**p*= or <0.001, and \*\**p*<0.01, with n.s. no significant difference).

Prior work demonstrates that ferrous Fe can displace Mg and thereby serve as a nanomolar inhibitor for multiple MST family enzymes, including isochorismate synthases (*Pseudomonas aeruginosa* PchA and *E. coli* EntC) and salicylate synthase (*Yersinia enterocolitica* Irp9) (32). We therefore hypothesized that Mn, previously shown to be inactive as a cofactor for isochorismate synthase (30), might serve as a general inhibitor of MST family enzymes in *B. subtilis*. To test this hypothesis, we asked whether supplemental Mg could suppress Mn sensitivity. Indeed, Mg reversed the Mn-dependent growth inhibition of the YS strain in both liquid culture (Fig 4c) and on plates (Fig 4d). Elevated levels of Mg also reversed the Mn-dependent growth inhibition of the PS and PSYS strains on LB medium (Fig. 4d). Thus, Mn-dependent inhibition of Mg-dependent enzymes is correlated with growth inhibition.

### Induction of MST family enzymes reverses Mn intoxication of YS but not PS

To test the hypothesis that inhibition of Mg-dependent MST enzymes is limiting for growth, we engineered strains with inducible expression of several of these enzymes. Surprisingly, induction of any of the three MST proteins suppressed Mn intoxication of the YS strain (Fig. 4e, f) but not the PS strain in LB medium (Fig. S3). Induction of these chorismate-utilizing enzymes had no effect on growth in the absence of Mn challenge (Fig. S4). These results suggest that Mn intoxication inhibits a similar set of enzymes in the PS and YS strains, but the specific metabolic deficiencies that underlie growth inhibition differ.

Of all the products of pathways dependent on MST enzymes, the simplest to assay is the siderophore bacillibactin and its precursors. In *B. subtilis* 168 strains this pathway is not fully functional due to the presence of the *sfp^0^* mutation that inactivates the Sfp 4-phosphopantetheinyl transferase necessary for the activity of non-ribosomal peptide synthases (34, 35). As a result, 168 strains fail to properly modify the DhbB and DhbF enzymes with the phosphopantetheine carrier and produce the bacillibactin precursors 2,3-dihydroxybenzoic acid (DHBA) and 2,3-dihydroxybenzoylglycine (DHBG), known collectively as DHBA(G) (36). These precursors are easily assayed in cell supernatants of iron-starved cells using spectrophotometry. We monitored DHBA(G) production using *fur* mutants that derepress the expression of the entire *dhb* operon for bacillibactin production (37). As expected, DHBA(G) production was elevated in cells grown in MM-malate in the *fur* mutant background. However, DHBA(G) production was not elevated if the medium was amended with Mn, or if the *fur* mutant was present in the YS mutant background (Fig. 4g, Fig. S5). Further, DHBA(G) production was increased even above the levels seen in the *fur* mutant in strains overexpressing the DhbC isochorismate synthase, and synthesis was no longer inhibited by Mn (Fig. 4g). DHBA(G) synthesis was also increased by expression of a different isochorismate synthase, MenF. These observations are consistent with the hypothesis that Mn is a generalized inhibitor of MST enzymes, and this inhibition can be overcome by Mg supplementation or by enzyme overproduction. Unexpectedly, the induction of *pabB* also increased the expression of DHBA(G) (Fig. 4g, Fig. S5). Although PabB is an MST-family enzyme, it is not known to have isochorismate synthase activity. Thus, the mechanism of this effect may be indirect, perhaps through the binding of Mn and protection of DhbC from Mn inhibition.

## DISCUSSION

Metalloenzymes are ubiquitous in biology, but their activity is contingent on metalation with the proper metal cofactor. Many metalloproteins obtain the cognate metal ion from a buffered cytosolic pool dependent on metal availability and the metal affinity of the protein active site (38). According to the Irving-Williams series, divalent transition metals typically bind with affinities in the order: Mn(II) < Fe(II) < Co(II) < Ni(II) < Cu(II) > Zn(II). These thermodynamic affinities often determine the hierarchy for metal occupancy within a specific metalloenzyme. The metalation of enzymes that require high-affinity metals, such as Cu or Zn, typically involves ligand exchange reactions with cellular metabolites or protein metallochaperones. For lower affinity metals, such as Mn and Fe, the buffered metal pools are more loosely held, and metal acquisition may occur unaided from pools of rapidly exchanging ions. In both cases, metalation by more tightly held and slower exchanging metals (from higher in the Irving-Williams series) can lead to enzyme mismetalation and inactivation. For example, Zn intoxication has often been linked to the mismetalation of proteins that normally require Mn or Fe for their activity (39–41).

Even though Mn and Fe are relatively weak-binding metals (at the lower affinity end of the Irving-Williams series), cells can still experience metal intoxication when their homeostasis systems are disrupted. In *B. subtilis*, Fe intoxication is apparent in strains lacking the PfeT Fe efflux ATPase (42, 43), and Mn intoxication is apparent in efflux-deficient PS strains (9, 11) and, as shown here, in the absence of the *yqgC-sodA* operon (YS). The molecular basis for intoxication by these more weakly interacting metal ions is not well understood, but previous results have suggested that excess Mn may interfere with Mg-dependent processes (14, 44). For example, mutations in a *B. subtilis* Mg-efflux system (MpfA) can increase tolerance to high Mn levels (14). However, the Mg-dependent pathways that are inhibited by Mn have not been defined. Prior work has shown that Mn intoxication of the PS mutant derives, in part, from a dysfunction associated with the QoxABCD terminal oxidase and is correlated with production of RRS (11).

Here, we focused on understanding the role of the *yqgC-sodA* operon in Mn homeostasis. The YS operon deletion mutant is phenotypically similar to the PS strain characterized previously (11). Both the YS and PS mutants display similar Mn sensitivity (Fig. 1d), accumulation of intracellular Mn when excess Mn is added (Table 1), Mn-induced production of RRS (Fig. 1e), and changes to their metabolomes (Fig. 4b, S2). In addition, the Mn sensitivity of both strains can be suppressed by high Mg levels (Fig. 4d). This similarity is supported by a lack of additivity between the PS and YS mutations (Fig. 1d), which is consistent with the observation that the YS strain is defective for expression of *mneP* and *mneS* (Fig. 2a), encoding two efflux proteins lacking in the PS strain. The mechanism underlying the reduced expression of *mneP* and *mneS*, and the lack of Mn induction (Fig. 2a), in the YS mutant is unclear but does not appear to be due to an inability of MntR to activate transcription (Fig. 2c).

Although the YS and PS strains are phenotypically similar, we were intrigued to observe that the YSPS strain accumulated less Mn than the YS strain after Mn shock (Table 1). One possibility is that the MneP and MneS CDF proteins contribute to Mn influx in the YS mutant strain under conditions of Mn excess. This is reminiscent of our previous observation that PS mutant strains lacking MntH, a proton-coupled Mn importer, accumulated more (rather than less) Mn when grown in Mn excess conditions (11). These findings lead to the hypothesis that these proton-coupled transporters may function in either direction, depending upon the strength of the relevant proton and metal ion gradients.

The phenotypes reported here are most evident in strains lacking both *yqgC* and *sodA*, but not in the single mutant strains. Superoxide dismutase is a broadly conserved protein found in all domains of life (45). In *B. subtilis*, MnSOD is the major Mn-binding protein in the cell (16) and protects cells against the reactive superoxide radical. The co-transcribed *yqgC* gene encodes an unknown function (DUF456) membrane protein. YqgC and its homologs (COG2839) are present in >1000 sequenced genomes, including *Deinococci*, *Mycobacteroides abscessus,* and *Salmonella enterica*. Although both *sodA* and *yqgC* are broadly conserved in many bacteria, gene neighborhood analysis reveals that the *yqgC*-*sodA* genomic organization is predominantly seen in the *Bacilli* class (Fig. S6).

It is not obvious why the deletion of both genes is required to reveal a high level of Mn sensitivity. However, this sensitivity is correlated with RRS accumulation, as reported previously (11). Our prior work linked RRS production to a dysfunction of the major quinol oxidase (QoxA)-dependent electron transport chain (ETC). One possibility is that the YqgC membrane protein interacts with proteins of the ETC or with menaquinone itself to help prevent RRS production, and in its absence MnSOD helps prevent RRS accumulation.

Both the YS and PS strains had similar metabolic profiles when challenged with Mn (Fig. S2). We noted that several of the over-represented metabolites are substrates of Mg-dependent enzymes, consistent with the finding that elevated Mg suppresses intoxication (Fig. 4d). However, the proximal cause of growth inhibition in these two strains appears to be different. Overexpression of MST family enzymes rescued the YS strain (Fig. 4e), but not the PS strain (Fig. S3). We therefore conclude that Mg-dependent MST family enzymes are inhibited in these strains, but other Mg-dependent processes also fail in the PS strain, precluding growth rescue by simply overexpressing MST enzymes. Expression of any of several MST enzymes can rescue growth (Fig. 4e), even though they do not all catalyze the same reaction (Fig. 4a). In some cases, the expressed enzyme may be directly acting to accelerate the limiting reaction, and in others, it perhaps acts as a decoy to help buffer toxic Mn ions in the cell. A similar effect may account for the ability of induction of PabB to help increase the expression of DHBA(G) (Fig. 4g).

Our findings here extend our view of Mn homeostasis in *B. subtilis* (Fig. 5). In addition to the core MntR-regulated import (*mntH*, *mntABC*) and efflux (*mneP*, *mneS*) genes, Mn is also trafficked in the cell by two TerC family efflux proteins in support of exoenzyme metalation (27). We here demonstrate that YqgC and MnSOD are two additional factors that have protective roles in Mn homeostasis. Finally, our results emphasize that there are likely multiple ways in which excess Mn can intoxicate cells. Even the phenotypically similar PS and YS strains appear to have distinct proximal causes for growth inhibition.

**Figure 5.**
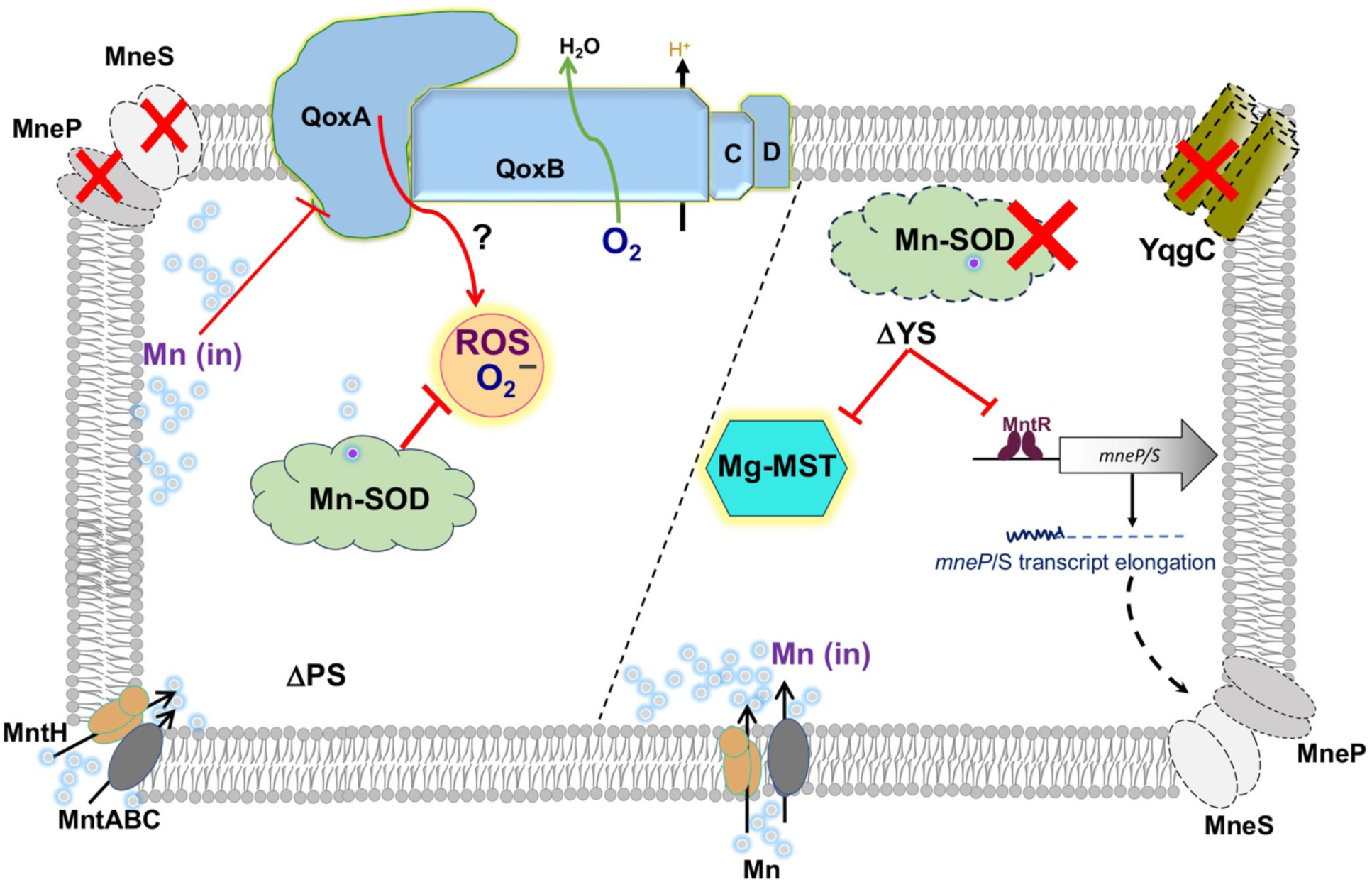
Functions of *yqgC*-*sodA* operon are needed for Mn homeostasis. In the absence of the MneP/S Mn efflux proteins cells accumulate Mn, which causes the production of ROS and other reactive radical species (RRS). This stress is reduced by deletion of quinol oxidase (Qox). Typically, cytosolic superoxide is eliminated by Mn-SOD encoded by *sodA*. The *sodA* gene is transcribed with *yqgC* encoding a DUF456 membrane protein. In the YS deletion strain, transcription of *mneP* and *mneS* is affected leading to Mn sensitivity. Mn intoxication inhibits Mg-dependent enzymes including MST enzymes. However, over-production of MST enzymes rescues YS cells, but not the PS strain, reinforcing the idea that the proximal cause of growth inhibition differs between these two phenotypically similar strains.

## Materials and Methods

### Bacterial strains and growth conditions

The strains used in this study are listed in Supplementary file Table S1A. Mutations from *B. subtilis* 168 were moved into *B. subtilis* CU1065 as the parental WT strain for this study. Cultures were streaked from frozen glycerol stocks onto LB agar plates and grown at 37 °C for 18 hours. Cells were grown in 5 ml of LB broth supplemented when required with antibiotics: MLS (1 μg/ml erythromycin plus 25 μg/ml lincomycin), kanamycin (15 μg/ml), and/or chloramphenicol (10 μg/ml), at 37 °C under shaking conditions of 300 rpm on a gyratory shaker. Once cultures reached an OD_600nm_ ∼0.4, 2 μl volume was used as an inoculum into 98 μl of LB dispensed in 96 well plates such that the initial OD_600nm_ was 0.07. These plates were supplemented with appropriate antibiotics, and IPTG/xylose was added as needed. Growth in LB or modified minimal media recipe (11) containing either glucose or malate supplemented with MnCl_2_ (Sigma) was monitored for up to 24 hr at 37 °C under shaking conditions using Synergy H1 (BioTek Instruments, Inc. VT) plate reader. The minimum inhibitory concentration (MIC) for Mn was defined as the Mn concentration that led to an OD_600nm_ of <0.4 after 8 hr of growth.

### Genetic manipulations and strain construction

**A) CRISPR editing** The complementation strain was constructed using a repair template consisting of upstream (of *yqgC*) and downstream (of *sodA*) homology regions fused to *yqgC*-S936-*sodA* central fragment amplified and stitched using primers listed in the Supplementary file Table S1B. DNA alleles were generated by adaptor-based LFH PCR and were subjected to stitching to generate a repair template. The repair template had compatible *SfiI* flanking the 5’ and 3’ ends. Upon digestion with *SfiI* at 50 °C for 2 hr, a similar restriction digest was performed for pAJS23 plasmid containing guide RNA against *erm* cassette as described previously (20). The ligation of the digested repair template into pre-digested pAJS23 was performed and was then transformed into *E. coli* DH5α selected on Luria-Bertani (LB) medium (Affymetrix) supplemented with 30 μg/ml of kanamycin. Plasmid was isolated and moved into *E. coli* TG1 strain. The multimeric plasmid was extracted and was then transformed into *yqgC*-S936-*sodA::erm* recipient strain grown to OD_600nm_ = ∼0.8 in the modified competence (MC) media. Plasmid DNA was used to transform recipient cells, which were then allowed to recover for 2 hr under aerobic incubation at 30 °C. Transformed *Bacillus* cells were selected on LB-kanamycin plus 0.2% mannose in the agar at 30 °C (until clones appeared, i.e., ∼48 hr). The CRISPR plasmid was cured at 45 °C for 2 days on LB agar with repeated passages for all transformants. Clones sensitive to kanamycin and erythromycin were selected, and chromosomal DNA was prepared for PCR analysis. The constructed strains were verified using Sanger sequencing of amplified PCR reactions. **B) Overexpression in pPL82 or pAX01**. For ectopic expression of *Bacillus* genes *(mneP, menF, dhbC, pabB, aroE, aroH, and menA)* and heterologous expression of *Staphylococcus (mntE)* and *E. coli (yiiP/fieF)* genes, the coding regions were amplified with primers listed in Supplementary file Table S1B from chromosomal DNA using Phusion high-fidelity DNA polymerase (NEB) and subjected to restriction enzyme digestion, purification, and ligation into pre-digested pPL82/pAX01 vector for propagation in *E. coli* DH5α on LB supplemented with 100 μg/ml of ampicillin. Prepared recombinant plasmid constructs were transformed into recipient *Bacillus* strains for insertion at the *amyE* or *lacA* locus by double-cross over recombination selected with 10 μg/ml chloramphenicol or 1 μg/ml of erythromycin plus 25 μg/ml of lincomycin). **C) Multiple (adjacent) gene deletions**. For *yqgC*-*S936*-*sodA* and *S936*-*sodA* deletions, upstream and downstream fragments were amplified with long flanking homology PCR and stitched together using joining PCR such that the donor fragment had *yqgB*-*erm*-*yqgE* (ΔYS) and *yqgC*-*cat*-*yqgE* (ΔS936-*sodA*). These fragments were then transformed into CU1065, and integrants were selected with antibiotics.

### Disk diffusion and Mn sensitivity assays

Mn sensitivity was evaluated on a minimal medium with malate or glucose as described (11) previously. Briefly, the bottom agar contained 15 ml MM-malate broth agar (with 1.5 % final agar concentrations), and the top soft-agar consisted of 4 ml MM-malate broth agar (with 0.75 % agar final agar concentrations) and 100 μl of mid-exponentially grown cultures (OD_600nm_ of 0.4) as inoculum. Whatman filter paper number 4 disks (6 mm in diameter) with Mn (10 μl of 10 mM stock) were placed on the plates, and the diameter for the zone of growth inhibition (ZOI) was measured after 18 hr at 37 °C.

### Real-Time RT PCR

Total RNA was extracted using a QIAGEN kit from 1.5 ml of mid-log (OD_600nm_ =0.4) WT and YS cells, which were grown in LB broth either in the presence or absence of 100 μM Mn. Total RNA was treated with DNase (Ambion) enzyme to further purify and remove traces of DNA. For each reaction, 2 µg of RNA was used for cDNA preparation using High-Capacity reverse transcriptase (Applied Biosystems) amplified with random hexamer primers. Further, for amplicon measurements 10 ng of cDNA was used as a template along with 500 nM of *mneP*, *mneS,* and *gyrA* (control) gene-specific qPCR F/R primers in a 1X SYBR green master mix (Bio-Rad). Threshold and baseline parameters were kept consistent for experiments performed on different days. All C_t_ mean values for the gene expression were normalized to *gyrA* (n=3).

### Mariner-transposon (mTn) mutagenesis

The YS cells containing pMarA plasmid were grown in LB broth with erythromycin (1 μg/ml) at 30 °C till OD_600_ reached ∼1.0, and cells were plated onto Mn gradient plate made up of MM containing glucose at 42 °C. Colonies that grew after overnight incubation near the high levels of Mn were sub-cultured in LB supplemented with kanamycin (15 μg/ml) at 37 °C to confirm the presence of mTn. Chromosomal DNA was purified from these ΔYS-Tn cells and was subjected to *TaqαΙ* restriction enzyme digestion at 37 °C for 2 h, followed by overnight ligation of cohesive ends to generate a circular transposon-gDNA chimeric library. The PCR reaction from ends of the mTn performed on the chimeric library was further subjected to Sanger’ sequencing analysis at Cornell BRC facility to identify the mTn insertion site within the genome (Supplemental Table S1D).

### Whole genome re-sequencing

Suppressors for YS strains were isolated using Mn-disk diffusion assay performed using either LB-with 150 μM Mn present in the broth, 100 μM Mn supplemented in a plate condition, or a disk diffusion performed in a MM-malate agar. For MM-malate agar, suppressors were picked from the clearance zone. All the suppressors were tested multiple times for the increased resistance to Mn and genomic DNA was extracted, purified, and subjected to Illumina sequencing (San Diego, CA) by MiGS or SeqCenter (Pittsburgh). The Illumina sequence reads were trimmed, mapped, aligned, and analyzed for sequence variants against the reference genome sequence NC_000964 using CLC genomics workbench software (Supplementary File Table S2). The nucleotide changes (relative to reference) found in both the parental WT (CU1065) and the YS (unstressed) strains were ignored, and only newly arising nucleotide variants found in the suppressor strains are indicated.

### Determination of intracellular metal content

ICP-ES analysis was performed using modifications of a previously described procedure (11). Briefly, strains were grown in 5 LB broth to an OD_600nm_ of 0.4 at 37 °C under aerobic growth conditions. Cultures were treated with or without 150 µM Mn and further grown for 30 min. 2 ml of samples were collected by centrifugation and washed twice with 5 ml Chelex-treated PBS, resuspended in 0.5 ml PBS, and weighed. Samples were digested in double distilled HNO_3_ in a carbon heat block using a Vulcan 84 automated sample digestion unit (http://www.qtechcorp.com/), followed by the addition of 60/40 nitric/perchloric acid and incubation at 150 °C. Samples were cooled down and resuspended in deionized 18 MOhm water with 5% HNO_3_. ICP analysis was run on an Agilent 7700 series inductively coupled argon plasma-optical emission spectrometer (ICP-ES) housed at the US Department of Agriculture—ARS Robert W. Holley Center for Agriculture and Health in Ithaca, NY (mean ± standard deviation [SD]; n=3 independent biological replicates, each containing three technical replicates).

### β-galactosidase plates and enzyme activity

All bacterial cultures were grown to a mid-log phase, streaked onto LB agar containing 100 μg/ml of X-gal and Mn (0-100 μM), and plates were incubated at 37 °C overnight. Blue color development for different strains was noted by capturing images.

### Metabolite extraction and measurements

All strains were grown in LB at 37 °C until the culture reached an OD600 of 0.4 and then treated with or without 0.15 mM (final concentration) of MnCl_2_ (Sigma-Aldrich) for 60 min at 37 °C. Cells were pelleted and quenched by resuspending in chilled 0.6 ml of mixtures of acetonitrile:methanol:water (40:40:20). These were further lysed using 0.1 mm Zirconia beads in a Precellys homogenizer (Bertin Instruments). Freshly lysed suspensions were centrifuged at ∼12,000 rpm for 8 min at 37 °C. The supernatants were passed through a 0.22 μm SpinX tube filter (Sigma-Aldrich). Extracted metabolites were separated on a Cogent Diamond Hydride Type C column of 1200 liquid chromatography coupled to an Mass 6220 TOF spectrophotometer (Agilent). Ion abundances of metabolites were estimated using Profinder 8.0 and log_2_ fold changes were analyzed and plotted using Gene Cluster 3.0 and Java Treeview.

### Quantification of Siderophores

*B. subtilis* cultures were aerobically grown overnight in MM-malate (1 μM of Fe) at 37 °C with and without 100 μM added Mn. Optical density at 600nm was noted, and cells were harvested by centrifugation at 5000 rpm for 10 min to retrieve supernatant, and to 1 mL supernatant 200 μl of 10 mM FeCl_3_ (Sigma) dissolved in 100 mM HCl was added. This supernatant fraction was then neutralized by the addition of 100 μl of 1 M Tris-HCl (pH 8.0), and the absorbance at 510 nm was recorded. A standard curve was prepared with DHBA (Sigma) under similar conditions to extrapolate levels of siderophores in unknown culture samples. All 510 nm values were normalized to culture OD at 600nm (n=3). Color development for different reaction tubes was captured using an iPhone XS (Apple, CA).

## Data availability statement

The metabolomics datasets generated during the current study are available in the Metabolomics Workbench (46) repository under accession number PR001779 and DOI http://dx.doi.org/10.21228/M8RM7K. All other data supporting the findings of this study are available within the paper and its supplementary information files.

## ACKNOWLEDGEMENTS

The authors acknowledge Shree Giri for conducting ICP-MS analyses, Eric Craft for the sample coordination, and Caitlin Leodis (’23) for technical assistance. This work was supported by National Institutes of Health grants R35GM122461 awarded to JDH and NIAID R25 AI140472 to KYR. The content is solely the responsibility of the authors and does not necessarily represent the official views of the National Institutes of Health.

## Author contributions

AJS, JDH project conception; AJS, VS, JJI, MP, YL performed experiments; AJS, VS analyzed data; KYR and JDH, funding acquisition and supervision; AJS, JDH drafting of manuscript; all authors helped edit and have approved final manuscript.

## REFERENCES

1. Chandrangsu P, Rensing C, Helmann JD. 2017. Metal homeostasis and resistance in bacteria. Nat Rev Microbiol 15:338–350.

2. Murdoch CC, Skaar EP. 2022. Nutritional immunity: the battle for nutrient metals at the host-pathogen interface. Nat Rev Microbiol 20:657–670.

3. Palmer LD, Skaar EP. 2016. Transition Metals and Virulence in Bacteria. Annu Rev Genet 50:67–91.

4. Becker KW, Skaar EP. 2014. Metal limitation and toxicity at the interface between host and pathogen. FEMS Microbiol Rev 38:1235–49.

5. Focarelli F, Giachino A, Waldron KJ. 2022. Copper microenvironments in the human body define patterns of copper adaptation in pathogenic bacteria. PLoS Pathog 18:e1010617.

6. Pi H, Patel SJ, Argüello JM, Helmann JD. 2016. The *Listeria monocytogenes* Fur-regulated virulence protein FrvA is an Fe(II) efflux P_1B4_ -type ATPase. Mol Microbiol 100:1066–79.

7. Neyrolles O, Wolschendorf F, Mitra A, Niederweis M. 2015. Mycobacteria, metals, and the macrophage. Immunol Rev 264:249–63.

8. Helmann JD. 2014. Specificity of metal sensing: iron and manganese homeostasis in *Bacillus subtilis*. J Biol Chem 289:28112–20.

9. Huang X, Shin JH, Pinochet-Barros A, Su TT, Helmann JD. 2017. *Bacillus subtilis* MntR coordinates the transcriptional regulation of manganese uptake and efflux systems. Mol Microbiol 103:253–268.

10. Que Q, Helmann JD. 2000. Manganese homeostasis in *Bacillus subtilis* is regulated by MntR, a bifunctional regulator related to the diphtheria toxin repressor family of proteins. Mol Microbiol 35:1454–68.

11. Sachla AJ, Luo Y, Helmann JD. 2021. Manganese impairs the QoxABCD terminal oxidase leading to respiration-associated toxicity. Mol Microbiol 116:729–742.

12. Fritsch VN, Loi VV, Busche T, Sommer A, Tedin K, Nürnberg DJ, Kalinowski J, Bernhardt J, Fulde M, Antelmann H. 2019. The MarR-Type Repressor MhqR Confers Quinone and Antimicrobial Resistance in *Staphylococcus aureus*. Antioxid Redox Signal 31:1235–1252.

13. Töwe S, Leelakriangsak M, Kobayashi K, Van Duy N, Hecker M, Zuber P, Antelmann H. 2007. The MarR-type repressor MhqR (YkvE) regulates multiple dioxygenases/glyoxalases and an azoreductase which confer resistance to 2-methylhydroquinone and catechol in *Bacillus subtilis*. Mol Microbiol 66:40–54.

14. Pi H, Wendel BM, Helmann JD. 2020. Dysregulation of Magnesium Transport Protects Bacillus subtilis against Manganese and Cobalt Intoxication. J Bacteriol 202.

15. Eymann C, Dreisbach A, Albrecht D, Bernhardt J, Becher D, Gentner S, Tam le T, Buttner K, Buurman G, Scharf C, Venz S, Volker U, Hecker M. 2004. A comprehensive proteome map of growing *Bacillus subtilis* cells. Proteomics 4:2849–76.

16. Tu WY, Pohl S, Gray J, Robinson NJ, Harwood CR, Waldron KJ. 2012. Cellular iron distribution in *Bacillus anthracis*. J Bacteriol 194:932–40.

17. Nicolas P, Mäder U, Dervyn E, Rochat T, Leduc A, Pigeonneau N, Bidnenko E, Marchadier E, Hoebeke M, Aymerich S, Becher D, Bisicchia P, Botella E, Delumeau O, Doherty G, Denham EL, Fogg MJ, Fromion V, Goelzer A, Hansen A, Härtig E, Harwood CR, Homuth G, Jarmer H, Jules M, Klipp E, Le Chat L, Lecointe F, Lewis P, Liebermeister W, March A, Mars RA, Nannapaneni P, Noone D, Pohl S, Rinn B, Rügheimer F, Sappa PK, Samson F, Schaffer M, Schwikowski B, Steil L, Stülke J, Wiegert T, Devine KM, Wilkinson AJ, van Dijl JM, Hecker M, Völker U, Bessières P, et al. 2012. Condition-dependent transcriptome reveals high-level regulatory architecture in *Bacillus subtilis*. Science 335:1103–6.

18. Irnov I, Sharma CM, Vogel J, Winkler WC. 2010. Identification of regulatory RNAs in *Bacillus subtilis*. Nucleic Acids Res 38:6637–51.

19. Koo BM, Kritikos G, Farelli JD, Todor H, Tong K, Kimsey H, Wapinski I, Galardini M, Cabal A, Peters JM, Hachmann AB, Rudner DZ, Allen KN, Typas A, Gross CA. 2017. Construction and Analysis of Two Genome-Scale Deletion Libraries for *Bacillus subtilis*. Cell Syst 4:291–305 e7.

20. Sachla AJ, Alfonso AJ, Helmann JD. 2021. A Simplified Method for CRISPR-Cas9 Engineering of *Bacillus subtilis*. Microbiol Spectr 9:e0075421.

21. Kalyanaraman B, Darley-Usmar V, Davies KJA, Dennery PA, Forman HJ, Grisham MB, Mann GE, Moore K, Roberts LJ, Ischiropoulos H. 2012. Measuring reactive oxygen and nitrogen species with fluorescent probes: challenges and limitations. Free Radical Biology and Medicine 52:1–6.

22. Murphy MP, Bayir H, Belousov V, Chang CJ, Davies KJA, Davies MJ, Dick TP, Finkel T, Forman HJ, Janssen-Heininger Y, Gems D, Kagan VE, Kalyanaraman B, Larsson NG, Milne GL, Nystrom T, Poulsen HE, Radi R, Van Remmen H, Schumacker PT, Thornalley PJ, Toyokuni S, Winterbourn CC, Yin H, Halliwell B. 2022. Guidelines for measuring reactive oxygen species and oxidative damage in cells and in vivo. Nat Metab 4:651–662.

23. Grunenwald CM, Choby JE, Juttukonda LJ, Beavers WN, Weiss A, Torres VJ, Skaar EP. 2019. Manganese Detoxification by MntE Is Critical for Resistance to Oxidative Stress and Virulence of *Staphylococcus aureus*. mBio 10.

24. Brown JB, Lee MA, Smith AT. 2021. Ins and Outs: Recent Advancements in Membrane Protein-Mediated Prokaryotic Ferrous Iron Transport. Biochemistry 60:3277–3291.

25. Ouyang A, Gasner KM, Neville SL, McDevitt CA, Frawley ER. 2022. MntP and YiiP Contribute to Manganese Efflux in *Salmonella enterica* Serovar Typhimurium under Conditions of Manganese Overload and Nitrosative Stress. Microbiology Spectrum 10:e01316–21.

26. Paruthiyil S, Pinochet-Barros A, Huang X, Helmann JD. 2020. *Bacillus subtilis* TerC Family Proteins Help Prevent Manganese Intoxication. J Bacteriol 202.

27. He B, Sachla AJ, Helmann JD. 2023. TerC Proteins Function During Protein Secretion to Metalate Exoenzymes. Nat Commun. 14:6186. doi: 10.1038/s41467-023-41896-1.

28. Bremer E, Calteau A, Danchin A, Harwood C, Helmann JD, Médigue C, Palsson BO, Sekowska A, Vallenet D, Zuniga A, Zuniga C. 2023. A model industrial workhorse: *Bacillus subtilis* strain 168 and its genome after a quarter of a century. Microb Biotechnol 16:1203–1231.

29. de Saizieu A, Vankan P, Vockler C, van Loon APGM. 1997. The *trp* RNA-binding attenuation protein (TRAP) regulates the steady-state levels of transcripts of the *Bacillus subtilis* folate operon. Microbiology 143:979–989.

30. Young IG, Gibson F. 1969. Regulation of the enzymes involved in the biosynthesis of 2,3-dihydroxybenzoic acid in *Aerobacter aerogenes* and *Escherichia coli*. Biochimica et Biophysica Acta (BBA) - General Subjects 177:401–411.

31. Hubrich F, Müller M, Andexer JN. 2021. Chorismate- and isochorismate converting enzymes: versatile catalysts acting on an important metabolic node. Chem Commun (Camb) 57:2441–2463.

32. Meneely KM, Sundlov JA, Gulick AM, Moran GR, Lamb AL. 2016. An Open and Shut Case: The Interaction of Magnesium with MST Enzymes. Journal of the American Chemical Society 138:9277–9293.

33. Lomelino CL, Andring JT, McKenna R, Kilberg MS. 2017. Asparagine synthetase: Function, structure, and role in disease. J Biol Chem 292:19952–19958.

34. Harwood CR, Mouillon J-M, Pohl S, Arnau J. 2018. Secondary metabolite production and the safety of industrially important members of the *Bacillus subtilis* group. FEMS Microbiology Reviews 42:721–738.

35. Ollinger J, Song KB, Antelmann H, Hecker M, Helmann JD. 2006. Role of the Fur regulon in iron transport in *Bacillus subtilis*. J Bacteriol 188:3664–73.

36. May JJ, Wendrich TM, Marahiel MA. 2001. The *dhb* operon of *Bacillus subtilis* encodes the biosynthetic template for the catecholic siderophore 2,3-dihydroxybenzoate-glycine-threonine trimeric ester bacillibactin. J Biol Chem 276:7209–17.

37. Pi H, Helmann JD. 2017. Sequential induction of Fur-regulated genes in response to iron limitation in *Bacillus subtilis*. Proc Natl Acad Sci U S A 114:12785–12790.

38. Osman D, Robinson NJ. 2023. Protein metalation in a nutshell. FEBS Lett 597:141–150.

39. Barwinska-Sendra A, Waldron KJ. 2017. The Role of Intermetal Competition and Mis-Metalation in Metal Toxicity. Adv Microb Physiol 70:315–379.

40. Foster AW, Young TR, Chivers PT, Robinson NJ. 2022. Protein metalation in biology. Curr Opin Chem Biol 66:102095.

41. Chandrangsu P, Helmann JD. 2016. Intracellular Zn(II) Intoxication Leads to Dysregulation of the PerR Regulon Resulting in Heme Toxicity in *Bacillus subtilis*. PLoS Genet 12:e1006515.

42. Guan G, Pinochet-Barros A, Gaballa A, Patel SJ, Argüello JM, Helmann JD. 2015. PfeT, a P_1B4_ -type ATPase, effluxes ferrous iron and protects *Bacillus subtilis* against iron intoxication. Mol Microbiol 98:787–803.

43. Pi H, Helmann JD. 2017. Ferrous iron efflux systems in bacteria. Metallomics 9:840–851.

44. Hohle TH, O’Brian MR. 2014. Magnesium-dependent processes are targets of bacterial manganese toxicity. Mol Microbiol 93:736–47.

45. Sendra KM, Barwinska-Sendra A, Mackenzie ES, Baslé A, Kehl-Fie TE, Waldron KJ. 2023. An ancient metalloenzyme evolves through metal preference modulation. Nat Ecol Evol doi:10.1038/s41559-023-02012-0.

46. Sud M, Fahy E, Cotter D, Azam K, Vadivelu I, Burant C, Edison A, Fiehn O, Higashi R, Nair KS, Sumner S, Subramaniam S. 2016. Metabolomics Workbench: An international repository for metabolomics data and metadata, metabolite standards, protocols, tutorials and training, and analysis tools. Nucleic Acids Res 44:D463–70.

47. Omasits U, Ahrens CH, Muller S, Wollscheid B. 2014. Protter: interactive protein feature visualization and integration with experimental proteomic data. Bioinformatics 30:884–6.

